# Derived loss of signal complexity and plasticity in a genus of weakly electric fish

**DOI:** 10.1101/2021.02.06.430027

**Authors:** David E. Saenz, Tingting Gu, Yue Ban, Kirk O. Winemiller, Michael R. Markham

## Abstract

Signal plasticity can maximize the usefulness of costly animal signals such as the electric organ discharges (EODs) of weakly electric fishes. Some species of the order Gymnotiformes rapidly alter their EOD amplitude and duration in response to circadian cues and social stimuli. How this plasticity is maintained across related species with different degrees of signal complexity is poorly understood. In one genus of weakly electric gymnotiform fish (*Brachyhypopomus*) only one species, *B. bennetti*, produces a monophasic signal while all other species emit complex biphasic or multiphasic EOD waveforms produced by two overlapping but asynchronous action potentials in each electric organ cell (electrocyte). One consequence of this signal complexity is the suppression of low-frequency signal content that is detectable by electroreceptive predators. In complex EODs, reduction of the EOD amplitude and duration during daytime inactivity can decrease both predation risk and the metabolic cost of EOD generation. We compared EOD plasticity and its underlying physiology in *Brachyhypopomus* focusing on *B. bennetti*. We found that *B. bennetti* exhibits minimal EOD plasticity, but that its electrocytes retained vestigial mechanisms of biphasic signaling and vestigial mechanisms for modulating the EOD amplitude. These results suggest that this species represents a transitional phenotypic state within a clade where signal complexity and plasticity were initially gained and then lost. We discuss potential the roles of signal mimicry, species recognition, and sexual selection in maintaining the monophasic EOD phenotype in the face of detection by electroreceptive predators.

## Introduction

Signal plasticity is one means for organisms to overcome trade-offs associated with costly communication signals. This is especially true for multi-functional signals such as echolocation calls or the electric organ discharges (EODs) of weakly electric fishes. Few studies have examined the evolution and maintenance of signal plasticity in a phylogenetic context. Here we compare signal plasticity in a genus of weakly electric fish and report a case in which a loss of signal complexity is associated with a strong reduction in signal plasticity, likely at great metabolic cost.

Gymnotiform fishes emit and detect weak electric fields to communicate with conspecifics and navigate in dark waters. The EOD is produced by the near-simultaneous action potentials of excitable cells (electrocytes) in the electric organ. Across their phylogeny, gymnotiforms have evolved diverse signals that vary in parameters such as discharge rate and waveform (Albert & Crampton 2005). The impressive diversity of EOD waveforms is due in part to differences in electrocyte morphology as well as the diversity, kinetics, and spatial distribution of the ion channels in these excitable cells (Markham 2013; Markham & Stoddard 2013). Despite its high species diversity and wide geographical range, the genus *Brachyhypopomus* (Hypopomidae) is well studied relative to other gymnotiforms, especially its phylogeny (Fig. 1) (Crampton et al. 2016a,b) and its ecology (Waddell et al. 2019), making these fishes particularly useful for comparative evolution and neurophysiology research.

**Figure 1.**
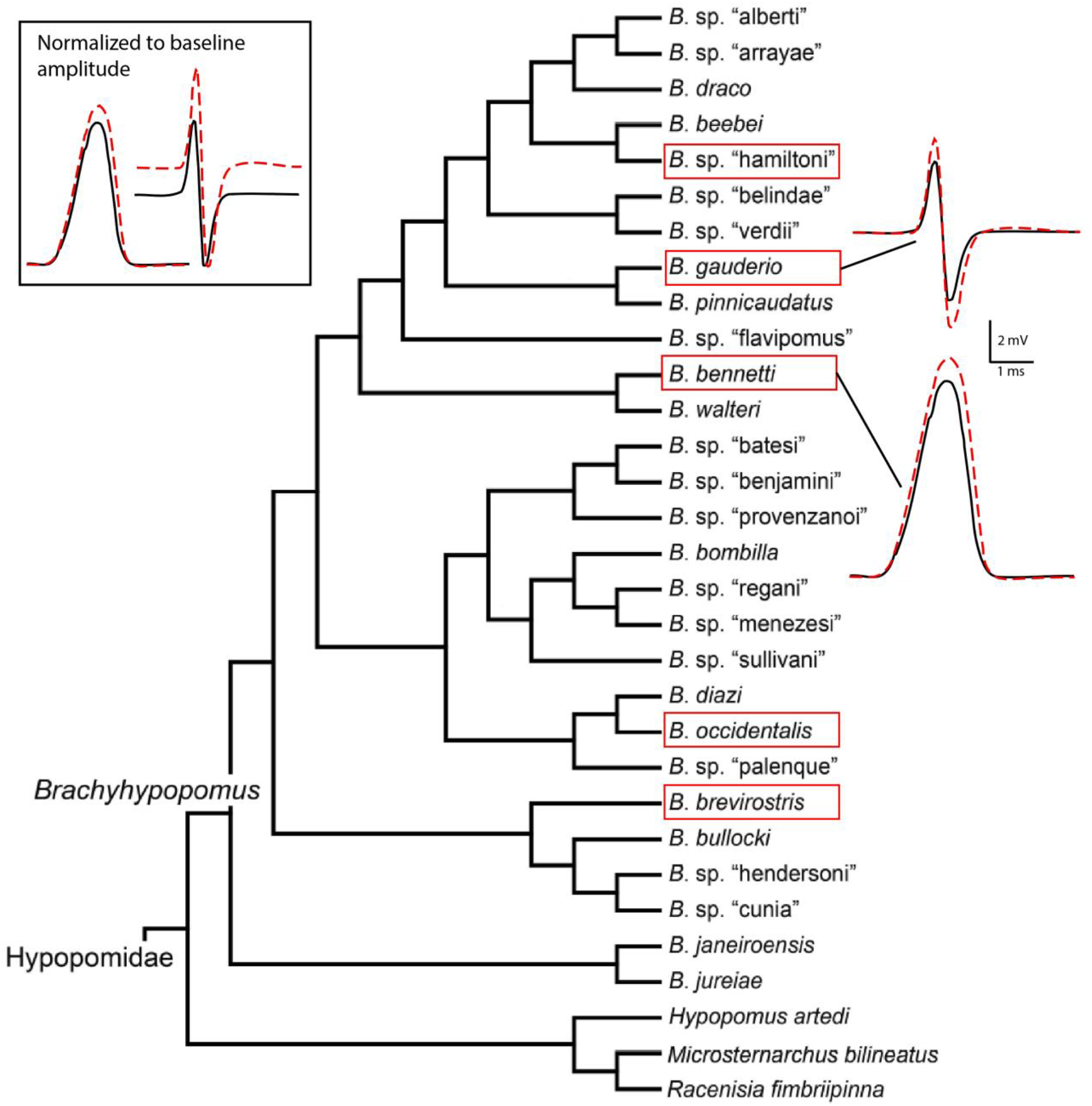
Current phylogeny of *Brachyhypopomus* adapted with permission from Crampton et al. (2016b). Species in red boxes are known to display circadian changes in EOD waveform. *B. occidentalis* was recorded by Hagedorn (1995). Insets at right show the EOD waveforms of the biphasic species *B. gauderio* and monophasic *B. bennetti*. Red dashed lines show nightly waveform changes. Inset at upper left shows the same EOD waveforms, with amplitudes scaled to the same baseline amplitude. The baseline EOD amplitude is larger in *B. bennetti*; however, the magnitude of the EOD amplitude change is much smaller.

Of the 28 *Brachyhypopomus* species described, the EODs of 18 species are known, 12 of which are biphasic, 5 are multiphasic, and 1 is monophasic (Fig. 1; Sullivan et al. 2013). Monophasy is a rare characteristic among weakly electric fishes, presumably because of its potential detection by electroreceptive predators (Stoddard 1999; Stoddard & Markham 2008). Of the larval *Brachyhypopomus* species recorded, all of them begin generating monophasic EODs and transition to more complex waveforms as they mature (Franchina 1997; Crampton et al. 2016b). This is thought to be the case for all gymnotiform fishes and the monophasic EOD is generally considered to be the ancestral trait (Stoddard 2002), though this has not been confirmed due to phylogenetic uncertainties between the families (Arnegard et al. 2010; Lovejoy et al. 2010; Tagliacollo et al. 2016; Alda et al. 2018).

In some gymnotiforms, crucial signal parameters such as phase, amplitude, duration, and low-frequency content vary by sex and ontogeny, and are regulated in response to seasonal, circadian, and social cues (Silva et al. 2002; Silva et al. 2007; Markham et al. 2009). This signal plasticity is regulated by steroid and peptide hormones, with peptide hormones playing a major role in regulating short-term circadian and social changes (Allee et al. 2009). Here we show how one form of short-term signaling plasticity that is regulated by melanocortin hormones varies between closely related species of *Brachyhypopomus*.

The general physiology of EOD is reviewed by Markham (2013; See Fig.1 therein) but, briefly, biphasic *Brachyhypopomus* electrocytes are large disc-shaped cells innervated on the posterior side (Bennett 1961, 1971; Trujillo-Cenoz et al. 1984). The biphasic EOD is the result of two action potentials (APs). The first AP (AP1) occurs at the posterior face of the electrocytes in the direction of the head and the second (AP2) occurs at the anterior face in the direction of the tail. The summation of the APs in a single electrocyte produces a biphasic single-cell discharge (μEOD) and the whole EOD is a sum of the composite μEODs

Previous work has shown that some gymnotiform species increase EOD amplitude (EODa) in response to circadian and social cues (Fig. 1), and that this amplitude plasticity is regulated by melanocortin hormones, such as adrenocorticotropic hormone (ACTH) (Markham & Stoddard 2005). Increasing EODa during nocturnal activity increases the metabolic cost as well as the active space of the signal, which carries implications for prey detection and for signal detection by both conspecifics and electroreceptive predators (Salazar & Stoddard 2008; Stoddard & Markham 2008; Stoddard et al. 2019). In *B. gauderio*, EODa is thought to be important for dominance displays and mate choice (Curtis & Stoddard 2003; Gavassa et al., 2012b).

All *Brachyhypopomus* species studied to date show pronounced modulations of EODa. To better understand the function and physiology of EODa plasticity, we compared the effects of ACTH in three additional species of *Brachyhypopomus*, and focused on one species in particular, *B. bennetti*, the only congener that emits a monophasic EOD. We expected that *B. bennetti* would show EODa plasticity on the same scale as other *Brachyhypopomus* species due to its relatively large EODa (Crampton & Albert 2006) and the potential conspicuousness of its monophasic EOD to electroreceptive predators (Stoddard 1999; Stoddard & Markham 2008). Additionally, we hypothesized that reversion to a monophasic EOD occurred in *B. bennetti* by the loss of voltage-gated Na^+^ channels and electrical excitability on the anterior face, matching the general organization of electrocytes in another gymnotiform with a monophasic EOD, the

Electric Eel (*Electrophorus electricus*) (Ellisman & Levinson 1982; Fritz et al. 1983). We found that *B. bennetti* exhibited only minor EODa plasticity and, more surprisingly, that the anterior membrane of *B. bennetti* electrocytes expresses voltage-gated Na^+^ channels and is electrically active.

## Materials and Methods

### Animals

A major limitation to working with this system is that it can be exceedingly difficult to reliably and responsibly acquire specific species. At this time, *B. gauderio* is the only species of *Brachyhypopomus* dependably bred in captivity. Individuals of *B. bennetti and B. brevirostris* were wild caught from Manaus, Brazil and exported in collaboration with researchers at the Brazilian Institute of Amazonian Research under ICMBio authorizations #14833 and #14834. These fish can be highly sensitive to the stressors of transport so ultimately we had only a total of 8 individuals of *B. bennetti* and 1 *B. brevirostris* for *in vitro* experiments. Specimens of *B. gauderio* were captive-bred from colonies maintained at the University of Oklahoma. All methods were approved in advance by the Institutional Animal Care and Use Committees of Texas A&M University and the University of Oklahoma.

### Immunohistochemistry

We immunolabeled voltage-gated Na^+^ channels and axon terminals in the electric organ using antibodies and protocols described previously (Ban et al., 2015). We labeled voltage-gated Na^+^ channels with an Anti-Pan Nav antibody (Alomone Labs, Jerusalem, Israel) that has been validated in electric fish (Ban, et al., 2015), and we labeled axon terminals with 3A10 (developed by T. Jessel and J. Dodd, and obtained from DSHB). For *B. bennetti tissue* (n=2 fish), selected sections were imaged on a Leica SP8 confocal laser scanning microscope using a 20x 0.75 NA oil objective. A GaN 405 laser with a 405 nm laser line was used to excite DAPI with an emission detection window between 415 nm and 455 nm. Similarly, an Argon laser line was used to excite the Alexa Fluor 488 and an emission detection window was set at 550 nm to 5500 nm. Z-series of images were acquired via sequential scanning. For *B. gauderio* tissues, immunolabeled sections were imaged on a Zeiss ApoTome.2 microscope with 5x/0.16NA, 10x/0.45NA, and 20x/0.80NA dry objectives. Images were acquired using a Zeiss AxioCam MRm and then processed by Zeiss AxioVision Rel.4.8. We created optical sections of the fluorescent samples using structured illumination. Image contrast was adjusted in Fiji and Adobe Photoshop for better visualization of fine structures in the electrocytes.

### Injections and recordings

EODs were recorded using a pair of silver electrodes on either side of the fish. EODs were amplified using a BMA 200 AC/DC Bioamplifier (CWE, Inc., USA) and digitized at 16 bits resolution using an A/D converter (CE Data translation USB Data Acquisition, USA) at a sampling rate of at least 50 kHz. Baseline EOD recordings were made for a minimum of 30 minutes. Fish were then quickly removed from the tank and given an intramuscular injection (1μl/g) of either 30 μM ACTH or normal saline in a process that took less than 15 seconds.

### Circadian recordings

Circadian recordings were conducted using an automated system for recording calibrated EODs round-the-clock from freely swimming fish described previously Stoddard et al. (2003). EODs were recorded at ~60-second intervals for at least three days to assess circadian variation in EOD waveform.

### Solutions for in vivo injections and in vitro electrophysiology

The normal saline contained 114 NaCl, 2 KCl, 4 CaCl_2_.2H_2_0, 2 MgCl_2_.6H20, 5 HEPES, 6 glucose; pH to 7.2 with NaOH. For some *in vitro* experiments with *B. bennetti* electrocytes, NaCl was substituted with choline chloride to reduce the Na^+^ concentration to 14.5 mM, and pancuroium bromide (1 μM) was added to the saline. Adrenocorticotropic hormone (ACTH from porcine pituitary) was purchased from Sigma Aldrich (St. Louis, MO)., Tetraethyl ammonium (TEA), barium chloride, and pancuronium bromide were obtained from Millipore-Sigma. Collagenase (type IV, Worthington Biochemical, Lakewood, NJ) was prepared in normal saline.

### Electrophysiology

To record discharges from single electrocytes (μEODs) we removed a small piece of the tail (~ 1cm) and carefully dissected the skin off to expose electric organ. The tissue was pinned into a Sylgard-coated recording dish and incubated in saline with 2% collagenase for 45 minutes to weaken the tissue surrounding the electrocytes. The preparation was then flushed several times with normal saline at RT (23 ± 1 °C) over a period of at least 15 minutes before recording.

### Current clamp

Intracellular stimulation and recordings used an Axoclamp 900A amplifier (Molecular Devices, Union City, CA) and extracellular recordings were completed using a Dagan TEV200A amplifier (Dagan Corp, Minneapolis, MN) in current clamp mode. A Digidata 1440 interface and PClamp 10.0 software (Molecular Devices, San Jose, CA) were used to control all amplifiers. Data were sampled and digitized at 100 kHz sampling rate. Extracellular pipettes were pulled from borosilicate glass and had resistances between 400-600 KΩ when filled with normal saline. Intracellular pipettes had resistances of 0.8-1.2 MΩ when filled with 3_M_ KCl.

We used a multi-electrode current clamp procedure to record APs from single electrocyte membranes in a procedure described in more detail elsewhere (Bennett 1961; Markham & Stoddard 2005, Markham & Zakon 2014). One intracellular pipette delivered a depolarizing current step to elicit the μEOD. A second intracellular pipette recorded the intracellular potential and two extracellular pipettes placed within 50 μm of the posterior and anterior membranes recorded extracellular potentials, one from each membrane. Off-line subtraction of the posterior extracellular record and the anterior extracellular record from the intracellular record result in AP1 and AP2 respectively. Subtraction of the posterior extracellular record from the anterior extracellular record returns the μEOD.

Only electrocytes with stable resting potentials < 80 mV and input resistances were recorded. Once all the electrodes were in place, we delivered 6-ms depolarizing current steps while manually adjusting the stimulus current magnitude until a μEOD was dependably elicited. A baseline recording was made of μEODs every 60 seconds for at least 15 minutes, after which we perfused normal saline for control cells or normal saline containing 100 nM ACTH. Due to the limited number of specimens, the timing of ACTH application and the length of the baseline recordings were staggered between individuals as an added measure of control for nonspecific effects. Solutions were changed during the interstimulus interval with a quick perfusion of 5 mL followed by slow continuous perfusion at 5 mL/h. We recorded μEODs at 60 second intervals for the remainder of the experiment.

### Two-electrode voltage clamp

Holding potential for all cells was −90 mV. Whole cell currents were recorded in normal saline in response to voltage steps from 140 mV to 0 mV in 5 mV increments. Na^+^ currents (I_Na_) were recorded in reduced-Na^+^ saline (14.5 mM NaCl) with 2 mM BaCl^2+^ and 10 mM TEA to block any K^+^ conductances, and 1μM pancuronium bromide to prevent spontaneous contraction of tail muscles. Voltage clamp protocols to assess Na^+^ current (INa) activation and inactivation consisted of a 50 ms conditioning step to −120 mV followed by 20 ms voltage steps from −120 to 25 mV in 5 mV increments and then a 20 ms step to 0 mV. Recovery of INa from inactivation was assessed with a protocol that consisted of a 50 ms conditioning step to −120 mV followed by an activation step to 0 mV and a recovery step to −120 mV for 0.5 ms to 12.5 ms in 0.5 ms increments and then a step to a test potential of 0 mV for 20 ms.

### Data treatment and analysis

For in vivo experiments, paired t-tests were used to compare percent changes in EOD parameters (relative to baseline) between in ACTH and saline. The same was done for the current clamp data. Statistical analyses were not performed for the injection data on *B*. cf. *hamiltoni* or for current clamp data on *B. brevirostris* because only one individual was available for each experiment (see results for modified procedure). Electrophysiology data were analyzed with Clampfit 10.7.0. (Molecular Devices) and Matlab (Mathworks, Natick, MA).

### Computational simulations

For numerical simulations we modeled the electrocyte as three cylindrical compartments arranged as a passive central compartment coupled to two flanking active compartments. External stimulation current was applied only to the central compartment, consistent with experimental procedure. The capacitance *CC* for the central compartment was 100 nF and the capacitances of the posterior and anterior compartments, C^P^ and C^A^ respectively, were 50 nF, yielding a total membrane capacitance of 150 nF consistent with empirical measurements of whole-cell capacitance. Differential equations were coded and integrated with Matlab using Euler’s method with integration time steps of 1 x 10^-9^ sec. All model parameters are shown in Table 1.

**Table 1.**
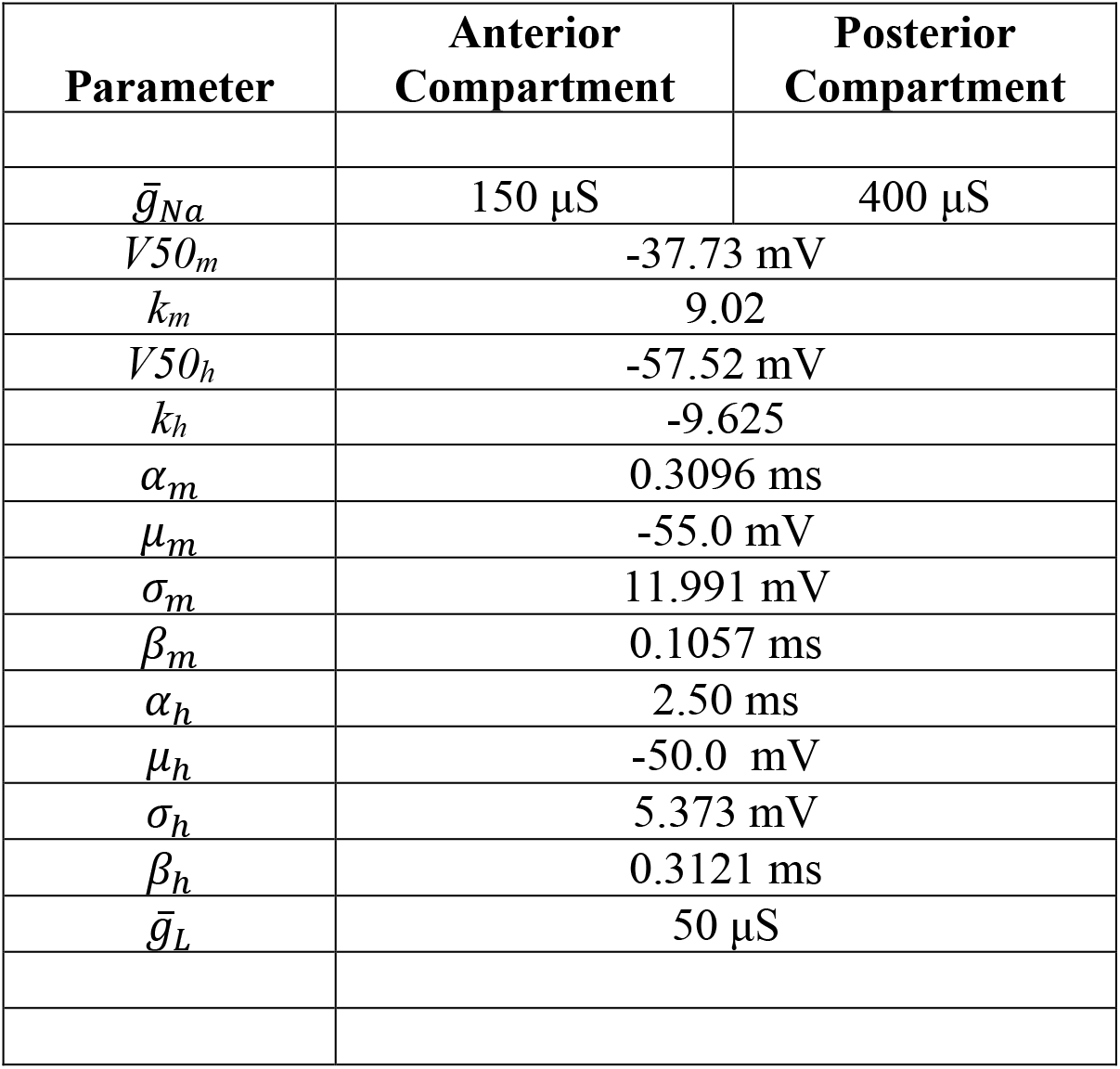
**Parameter values for the electrocyte model**.

| Parameter | Anterior Compartment | Posterior Compartment |
| --- | --- | --- |
| $\bar{g}_{Na}$ | 150 μS | 400 μS |
| $V50_m$ | -37.73 mV | |
| $k_m$ | 9.02 | |
| $V50_h$ | -57.52 mV | |
| $k_h$ | -9.625 | |
| $\alpha_m$ | 0.3096 ms | |
| $\mu_m$ | -55.0 mV | |
| $\sigma_m$ | 11.991 mV | |
| $\beta_m$ | 0.1057 ms | |
| $\alpha_h$ | 2.50 ms | |
| $\mu_h$ | -50.0 mV | |
| $\sigma_h$ | 5.373 mV | |
| $\beta_h$ | 0.3121 ms | |
| $\bar{g}_L$ | 50 μS | |

The passive central compartment’s current balance equation included only terms representing the injected stimulation current pulses I_Stim_(*t*), passive leak (I_LC_), and coupling to the two active compartments:

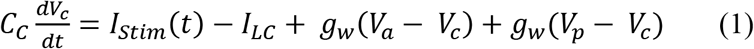

where *g_w_* is the coupling conductance, fixed at 7 μS for all compartments, *s* is the coupling ratio, fixed at 0.334, V is the membrane potential, and I_LC_ is the passive leak current given by the equation

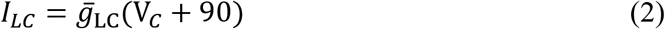

where *ḡ_LC_* = 5 μS.

The current balance equations for the anterior and posterior active compartments were, respectively:

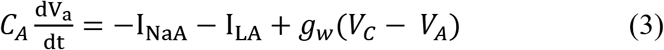

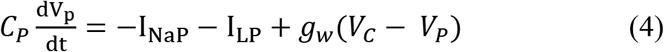

where V is the membrane potential, I_Na_ represents Na^+^ current, and I_L_ is the leak current. Equations for these currents were as follows:

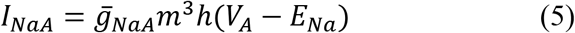

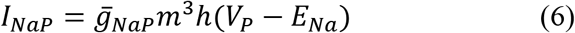

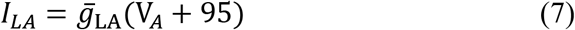

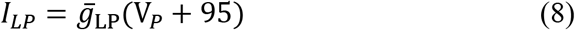

The gating variables in Equations 5 and 6 are evolved by Equation 9 where j = m or h:

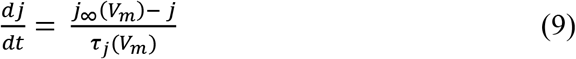

The values of *j*_∞_ were determined in a voltage-dependent fashion as follows:

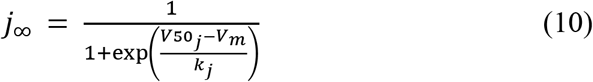

where *V*50_j_ and *k_j_* are derived from Boltzmann sigmoidal fits to experimental data and are given in Table 1. τ_j_ is given by Eqn. 11 for j = m:

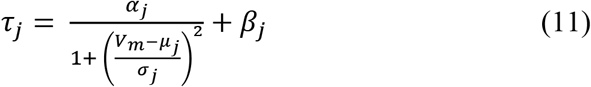

and by Eqn. 12 for j = h:

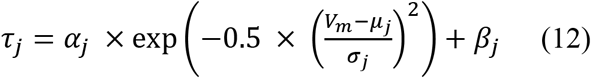

Where values of α*_j_*, β*_j_*, μ*_j_*, and σ*_j_* were determined by least-squares best fits to our experimental voltage-clamp data and are given in Table 1.

## Results

### Immunolocalization

Immunohistochemistry confirmed that electrocytes are innervated on the posterior stalk by spinal electromotor neurons in *B. bennetti* and *B. gauderio* (Fig. 2G-L and Fig. 3D-E, respectively). As expected in *B. gauderio*, Na^+^ channels were found on both the posterior and anterior membranes (Fig. 3A-C). Contrary to our expectations, Na^+^ channels were also found on both the posterior and anterior membranes of *B. bennetti* (Fig. 2D-F and Fig. 4). Interestingly, Na^+^ channels also appeared on internal cell structures (Fig. 3B, 4F). Due to limited availability of specimens, we were unable to quantify the Na^+^ channels. *B. bennetti* electrocytes are large (~600-900 μm in length) and the labeling of Na^+^ channels varies depending on where the cells were sectioned.

**Figure 2.**
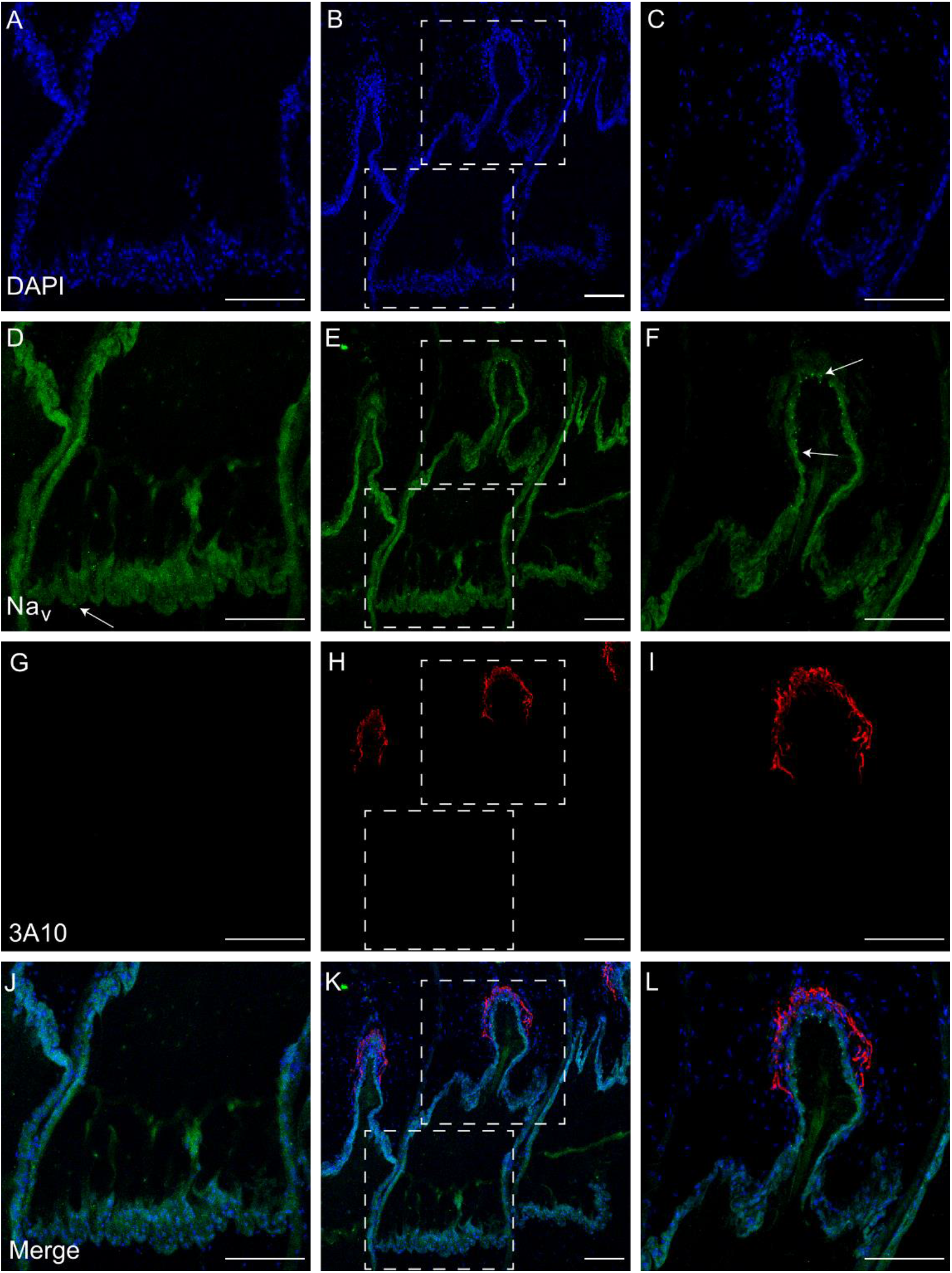
Immunolabeling of axon terminal and voltage-gated Na^+^ channels in electrocytes of the monophasic *B. bennetti*. The center column shows the full cell. Dashed boxes show delineate enhanced images of the anterior membrane shown on the left column and the posterior membrane shown on the right. (A-C) DAPI labels the electrocyte nuclei and provides a relative outline of electrocytes. (D-F) Anti-Pan NaV labels Na^+^ channels. Punctuated dots are visible on both membranes as well as on internal structures. (G-I) 3A10 labels the axon terminal on the stalk. (J-L) Merged images of the three previous rows. Scale bar: 100 μm

**Figure 3.**
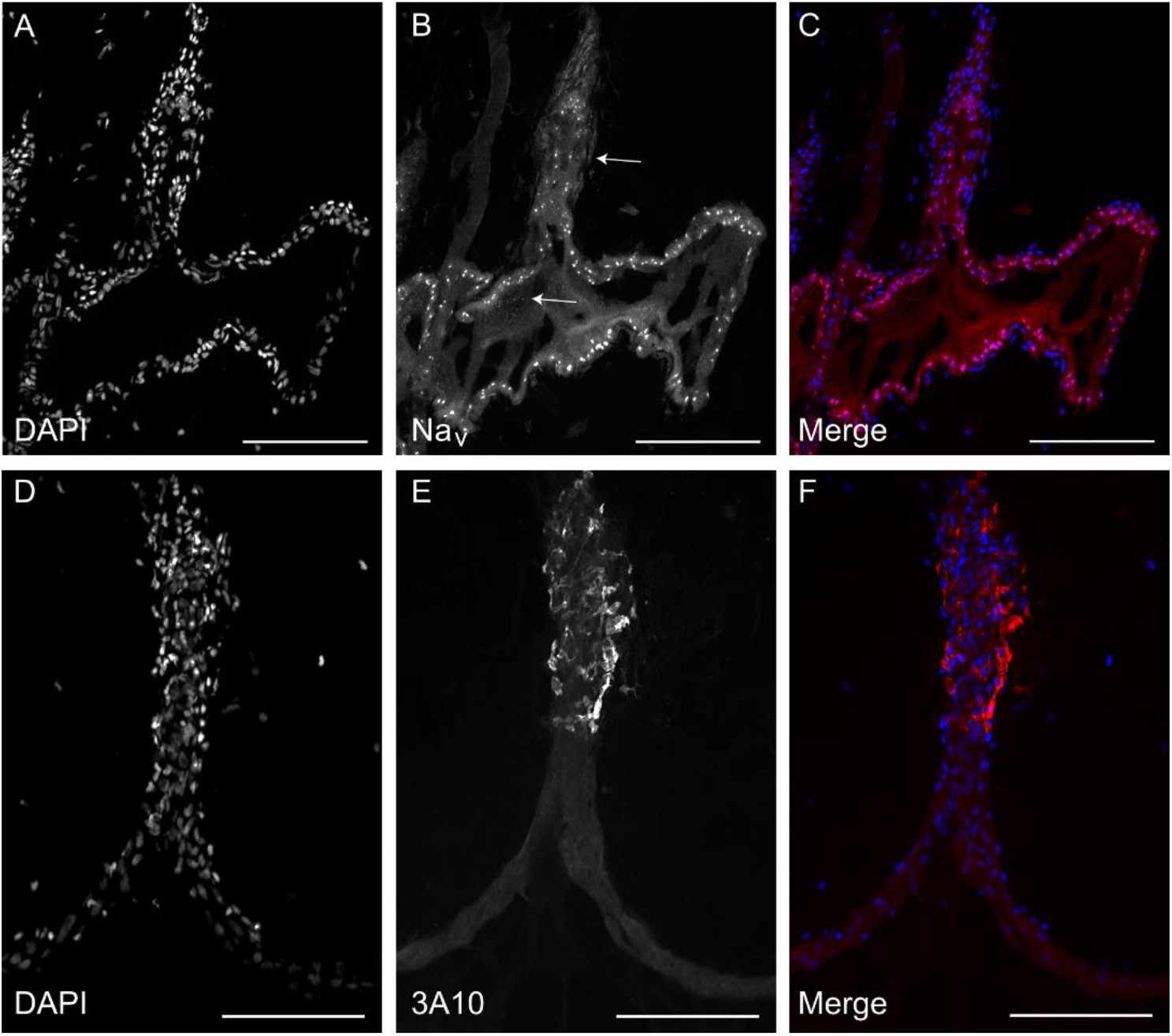
Immunolabeling of axon terminal and voltage-gated Na^+^ channels in the electrocytes of the biphasic species *B. gauderio*. A) DAPI labeling of nuclei provides the general outline of the electrocyte. B) Anti-Pan NaV labels Na^+^ channels. The central arrow highlights puncta present on internal structures. C) A & B merged. D) DAPI labels the nuclei outlining the stalk of a different cell. E) 3A10 labels the axon terminal on the stalk. F) D & E merged. Scale bar: 100 μm

**Figure 4.**
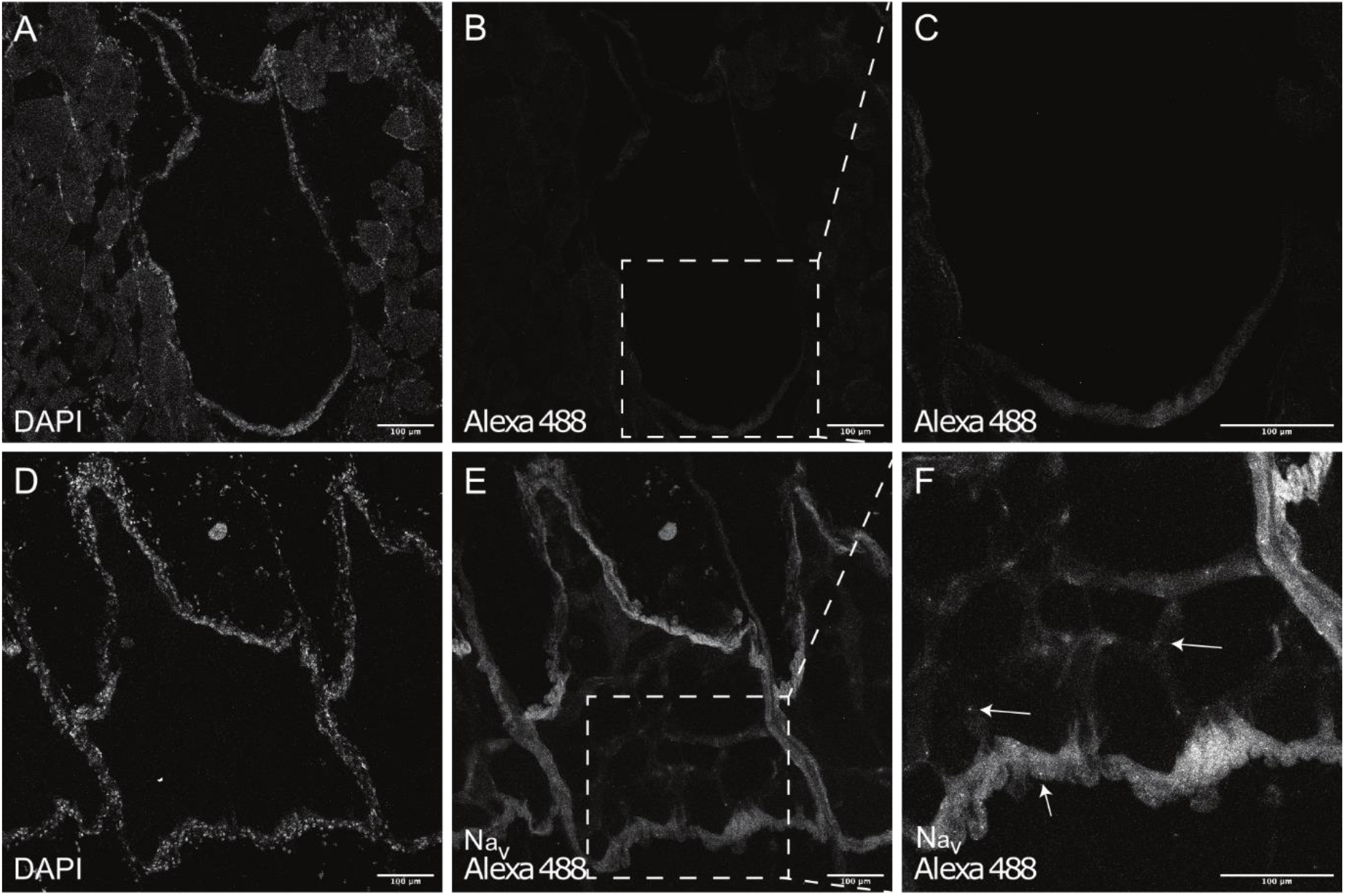
Na^+^ channels were present in both anterior (up) and posterior (down) membranes of the electrocytes in the monophasic species *B. bennetti*. (A and D) DAPI labels the electrocyte nuclei and provides a relative outline of electrocytes with the caudal stalk prominently visible. Control cells with only the secondary antibody (Alexa Fluor 488; B and C) show minimal autofluorescence but no positive signals indicating the specific signals (E and F) come from primary antibody (Anti-Pan Nav) and the presence of voltage-gated Na^+^ channels in the electrocyte membrane. High magnification image revealed the punctuated dot signals (arrows) located on the anterior membrane, contrary to our expectations (F). Punctuated dots are also visible on internal structures of electrocytes (F) Scale bar: 100 μm

### Effects of ACTH in vivo

We found that intramuscular injections of ACTH significantly increased the peak-to-peak EODa in the biphasic species, *B. brevirostris*, and *B*. cf*. hamiltoni* (33.7 ± 4.7% and 16.1%; n=6 and 1, respectively), as has been shown for *B. gauderio* (Markham & Stoddard, 2013). Separate evaluation of the phases in *B. brevirostris* showed that amplitude of both P1 and P2 increased (t=2.23, p = 0.056). On average, ACTH also increased EODa in *B. bennetti*, however, the effect was considerably weaker (t = 2.45, p = 0.094; n=6). Saline controls slightly decreased the EODa in all four species. The duration of both P1 and P2 (measured at 2% of the peak amplitude) increased in *B. brevirostris* (p < 0.001, t = 2.57) and *B*. cf*. hamiltoni*. In *B. bennetti*, the duration of the EOD did not change appreciably.

### Effects of ACTH in vitro

Using a multi-electrode current clamp, we measured μEOD parameters from individual electrocytes of *B. brevirostris* and *B. bennetti*. As previous studies have shown in *B. gauderio* (Markham & Stoddard 2005; Markham et al. 2009; Markham et al. 2013), ACTH increased amplitudes of both P1 and P2 of the μEOD in *B. brevirostris*, and concurrently, the total μEOD amplitude (Fig. 5C-D), the duration of P2, and the AP1-AP2 delay (data not shown). Because only one individual was available for testing, we attempted to wash out the ACTH after 30 minutes with saline (60 min of the total recording), during which we observed a slow decrease in μEOD amplitude over the course of the next hour. We then reapplied ACTH and saw a small increase again in the μEOD amplitude (Fig. 5D). In *B. bennetti*, ACTH also increased the amplitude of the μEOD (Fig. 5A-B; n=4), however the effect was much smaller relative to *B. gauderio* and *B. brevirostris*. Again, contrary our expectations, we observed APs on both the posterior and the anterior membrane (Fig.5A). Perfusion of ACTH slightly increased both AP1 and AP2. No other parameters showed consistent changes in response to ACTH.

**Figure 5.**
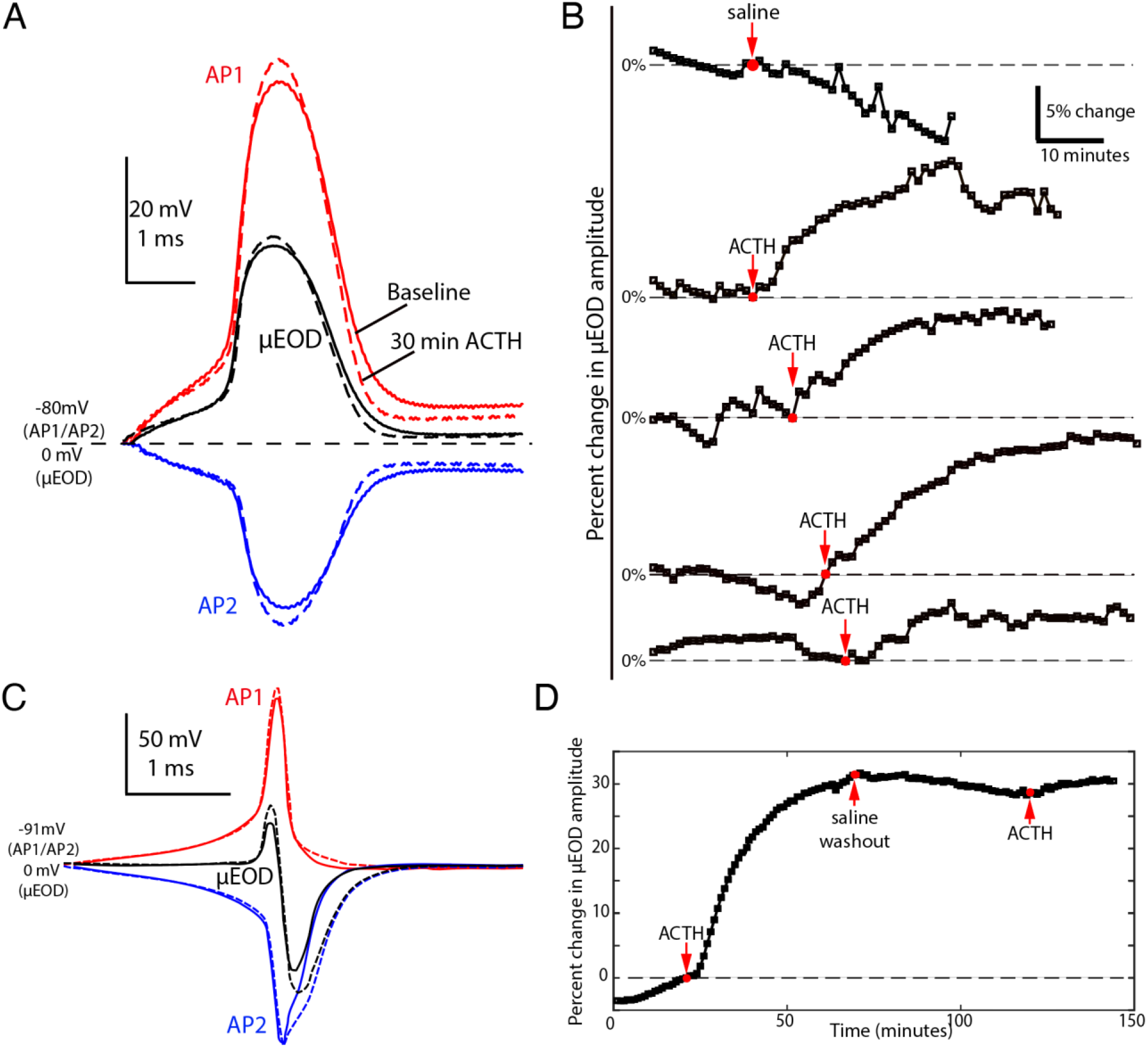
Electrocyte action potentials and response to ACTH in *B. bennetti* and *B. brevirostris* electrocytes. **A**, Representative recordings from a *B. bennetti* electrocyte of AP1 (red) and AP2 (blue) with the resulting μEOD (black). Dashed lines show the action potentials and μEOD 30 minutes after perfusion of ACTH. **B**, Recordings showing percent change in μEOD amplitude for 5 different *B. bennetti* individuals, after perfusion of either a saline control (1) or ACTH (4). **C**, Recordings from a *B. brevirostris* electrocyte of AP1 (red) and AP2 (blue) with the resulting μEOD (black). Dashed lines show the action potentials and μEOD 30 minutes after perfusion of ACTH. **D**, Percent change in μEOD amplitude for one B. brevirostris electrocyte in response to ACTH.

### Ionic currents in Brachyhypopomus bennetti electrocytes

Whole-cell currents recorded in voltage clamp showed an extremely large transient inward current that could not be controlled by the amplifier even at its maximum gain (50,000 V/V). Currents at steady state could be controlled by the amplifier’s steady-state gain (10^6^ V/V). Steady state I-V relationships were linear, showing no evidence of meaningful inward- or outward-rectifying K^+^ conductances (Fig. 6A).

**Figure 6.**
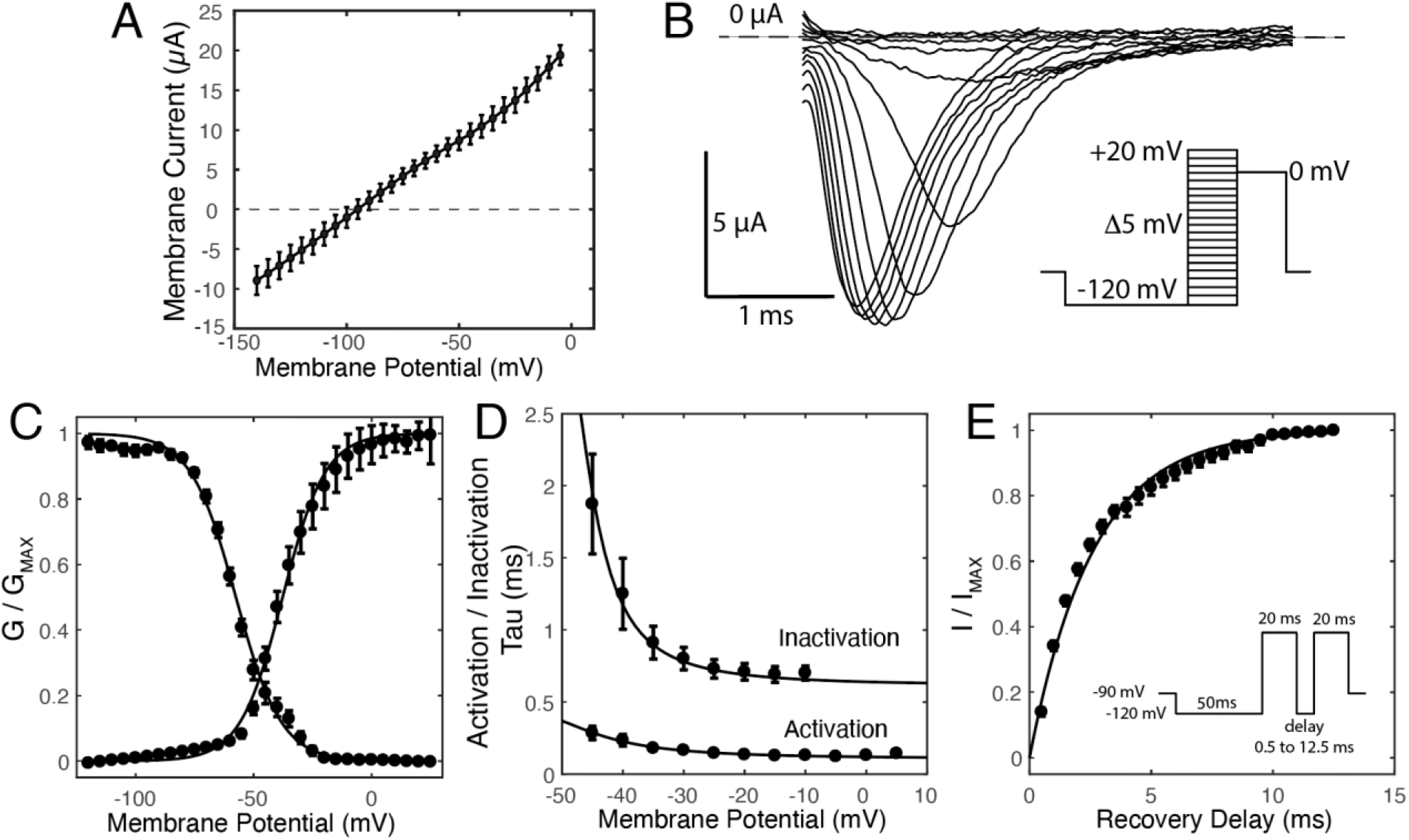
Characterization of *B. bennetti* Na^+^ currents. **A)** Whole-cell I-V relationships at steady state in *B. bennetti* electrocytes are linear and show no evidence of significant inward- or outward-rectifying K^+^ currents. In all panels symbols are means and error bars represent SEM. **B)** Representative currents elicited by voltage steps from −115 to 20 mV in 5 mV increments. Inset is voltage command protocol, which was also used for data shown in C and D. **C)** Activation and inactivation GV plots for Na^+^ currents. Solid lines are Boltzmann sigmoidal fits. **D)** Activation and inactivation τ-V curves. Solid line is a Gaussian fit for inactivation data and a Lorentzian fit for activation data. **E)** I_Na_ recovery from inactivation. Solid line represents a single-exponential fit. Inset is the voltage command protocol.

Given the apparent absence of significant voltage-gated K^+^ currents and the paucity of subject animals, we focused on describing the Na^+^ conductances in *B. bennetti* electrocytes because they are apparently the dominant ionic conductance shaping the electrocyte APs. We used a reduced-Na^+^ extracellular saline solution (14.5 mM NaCl) to reduce the driving force on I_Na_ and allow stable voltage clamp control during I_Na_ activation. *B. bennetti* electrocytes showed transient inward Na^+^ currents (Fig. 6B-C) with typical voltage dependence of activation and inactivation time constants (Figure 6D). The time constant of recovery from inactivation (τ = 3.80 ms; Fig. 6E) is the slowest for any gymnotiform species thus far reported, by a factor of four (cf. Ferrari et al., 1995; Markham et al., 2013; Markham and Zakon, 2014). This is not completely unexpected considering *B. bennetti* has a very low EOD frequency (<10 Hz).

### Computational electrocyte simulations

With parameters derived from our voltage clamp data, we simulated *B. bennetti* electrocytes to evaluate whether our experimentally-observed Na^+^ channel localizations, Na^+^ conductances, and electrocyte passive properties are sufficient and necessary to reproduce the electrocyte APs recorded *in vitro*, and to reproduce the effects of ACTH observed during our current clamp recordings. The model electrocyte, which included Na^+^ conductances in both the posterior and anterior compartments, produced simulated AP1, AP2, and μEOD waveforms (Fig. 7A) that were strikingly consistent with experimentally recorded waveforms (cf. Fig. 5A). Increasing the Na^+^ conductance by 10% in both anterior and posterior compartments increased AP1, AP2, and μEOD amplitude in the model cell (Fig. 7A) in a manner nearly identical to the effects of ACTH recorded *in vitro* (cf. Fig. 5A).

**Figure 7.**
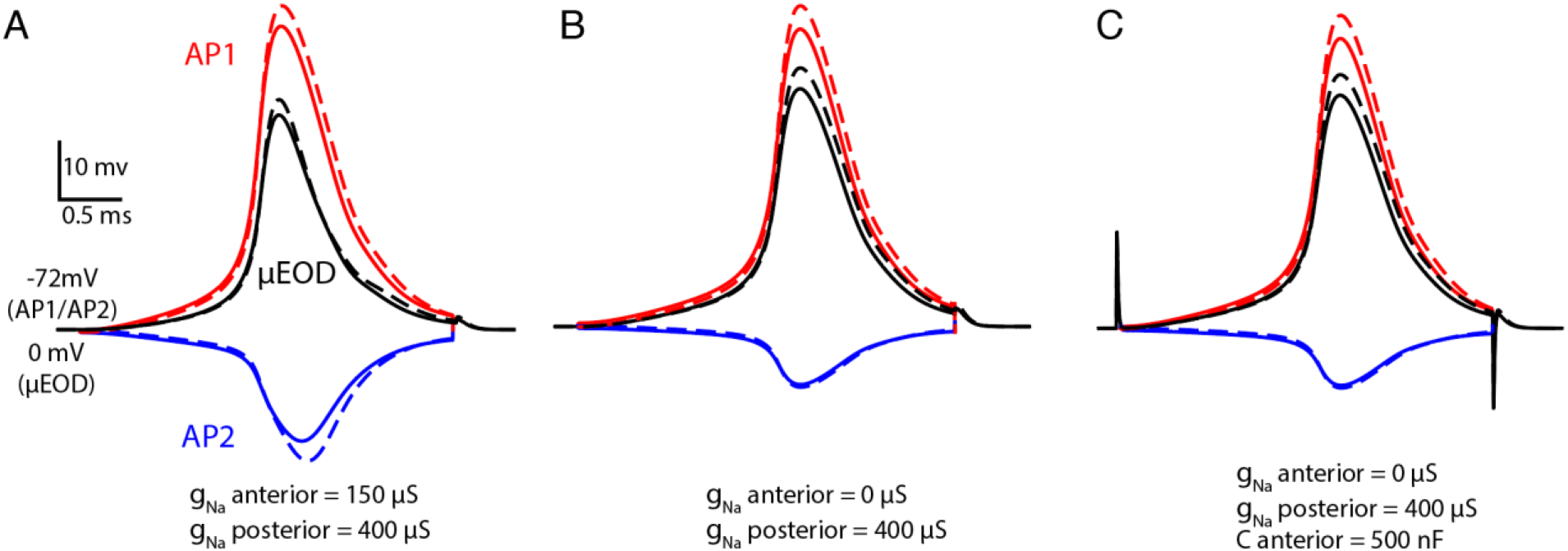
Computational simulations of *B. bennetti* electrocytes. Simulation of action potentials based on experimental measurements of Na^+^ conductance parameters under voltage clamp. Stimulation is a 3-ms step current of 4,400 nA (not shown). Passive membrane responses to stimulus current are not shown. Red lines represent the posterior compartment, blue lines represent the anterior compartment, and black lines represent whole-cell μEODs. Solid lines represent baseline simulations and dashed lines show simulations after Na^+^ conductances are increased by 10% in both the anterior and posterior compartments. Scale bars apply to all figure panels. **A)** Responses of baseline model electrocyte with all parameters as in Table 1. Action potentials closely resemble those recorded experimentally (compare to Fig. 5A). **B)** Responses of a model electrocyte where the anterior Na^+^ conductance is eliminated. **C)** Responses of a model electrocyte where the anterior Na^+^ conductance is eliminated and the anterior compartment capacitance is increased by a factor of 10 to a value of 500 nF.

To test whether Na^+^ conductances in the anterior compartment are necessary to reproduce our experimental results, we evaluated the performance of two alternate model electrocytes. In one model, we eliminated the anterior-compartment Na^+^ conductance. This model failed to reproduce experimentally-recorded AP1, AP2, and μEOD waveforms, as well as the observed effects of ACTH (Fig. 7B), even across a broad range of posterior-compartment Na^+^ conductance densities (data not shown). In a second alternate model, we eliminated the anterior-compartment Na^+^ conductance, but increased the anterior-compartment capacitance up to tenfold in order to test an earlier proposal that AP2 is a result of capacitative discharge from the anterior membrane (Bennett, 1971). This model also failed to reproduce experimental findings (Fig. 7C) even across a broad range of anterior-compartment capacitances and posterior-compartment Na^+^ conductance densities (data not shown). Taken together with our experimental data, these simulations support the conclusion that both anterior and posterior compartments of *B. bennetti* electrocytes are electrically active, and endowed with voltage-gated Na^+^ channels.

## Discussion

An obvious limitation of the experiments reported here is the limited sample size, a sometimes unavoidable consequence of collecting specimens in the wild. A further difficulty when working with specific species of gymnotiforms is that they are rarely, if ever, available through commercial fish importers because these fish can be tremendously difficult to identify at the species level without conducting detailed morphological analysis or EOD recordings. We compensated for restricted sample sizes wherever possible by using staggered baselines and other such strategies as described in the methods and results. Nonetheless, caution is called for with respect to the most striking observations here: that *B. bennetti* shows greatly reduced EOD plasticity compared to other Brachyhypopomid species and that both the anterior and posterior membranes of *B. bennetti* electrocytes are electrically active. In support of these conclusions, however, it is important to note that multiple approaches and sources of evidence (*in vivo* recordings, *in vitro* electrophysiology, immunolabeling, and computational simulations) all converged on the same outcomes.

### EOD plasticity in Brachyhypopomus

Relative to the other congeners studied, *B. bennetti* exhibited minimal EOD plasticity. Previous studies have shown that *B. gauderio* significantly modifies its EOD waveform in response to circadian and social cues via direct action of melanocortin peptide hormones on the electrocytes (Markham & Stoddard 2005; Markham & Stoddard 2013). Circadian plasticity of the EOD waveform has also been reported in *B. occidentalis* (Hagedorn 1995). Here we report similar ACTH-mediated EOD plasticity in three additional species of *Brachyhypopomus (B. bennetti, B. brevirostris* and *B*. cf. *hamiltoni*).

The extent of this EOD plasticity and its regulation varies by sex and ontogeny in *B. gauderio* (Markham & Stoddard 2013). Though our sample size is insufficient to confirm demographic variation for the species tested here, it is interesting that the plasticity seems consistent throughout the genus (Fig. 1) with *B. bennetti* as an exception. While some fishes from other gymnotiform genera exhibit ACTH-mediated EODa plasticity (McAnelly & Zakon 1996; Markham et al 2009), others do not (Saenz, unpublished data). The function of this plasticity is not yet fully understood, but two probable roles are social and energetic. The active electrosensory system can be metabolically costly. Estimates of EOD production costs range from 4% to 22% of the daily metabolic budget in species with relatively slow EOD repetition rates such as *B. gauderio* (Salazar & Stoddard 2008) and may cost well over 30% in species with higher repetition rates such as *Eigenmannia virescens* (Sternopygidae) (Lewis et al. 2014). Previous studies have reasonably argued that circadian regulation of EODa is adaptive for conserving energy. Therefore, it is interesting that *B. bennetti* shows decreased EODa plasticity, especially considering that its EODa is 3 to 8 times larger than those of sympatric congeners (Crampton & Albert 2006) and the present results suggest that the physiology of EOD production is energetically inefficient in this species.

Another surprising finding was the magnitude of AP2 in *B. bennetti* electrocytes and the presence of voltage-gated Na^+^ channels on the anterior membrane. The current clamp data suggest that, while the EOD of *B. bennetti* is monophasic, the μEOD is still composed of two APs. Because these APs completely overlap, the discharge from the anterior face (AP2) negates a significant fraction of AP1, potentially wasting considerable energy. Bennett (1971) recorded electrocytes from a monophasic Hypopomid, likely *B. bennetti*, but argued that the anterior membrane is electrically inexcitable. Instead, it was suggested that a smaller spike from the uninnervated membrane is a discharge of the membrane’s capacitance, similar to what is suggested to occur in the uninnervated face of electrocytes in *Gymnarchus niloticus*, an African weakly electric fish (Bennett 1971; Schwartz et al. 1975). Our finding that voltage-gated Na^+^ channels are indeed present on the anterior face, in conjunction with our electrophysiology data and computational simulations, suggests the anterior membrane is, in fact, excitable.

The presence of Na^+^ channels and generation of AP2 on the anterior membrane provides evidence against the long-standing assumption that *B. bennetti* s monophasic EOD is a retention of the pedomorphic monophasic condition. Is this electrical excitability of the anterior membrane a costly vestige of *B. bennetti* s biphasic ancestry or does AP2 serve some unknown function? While the sample sizes of the *in vitro* assays are admittedly small, it is nonetheless compelling that multiple lines of evidence from each assay point to the same conclusion: *B. bennetti* electrocytes have Na^+^ channels on the anterior membrane that contribute to a second membrane depolarization of significant magnitude. Consequently, AP2 decreases the energetic efficiency of the EOD, without the benefit of producing a biphasic EOD to cloak the EOD from electroreceptive predators.

### The persistence of the monophasic EOD

The biphasic EOD is likely an adaptation to predation pressures from eavesdropping electroreceptive predators (e.g. catfishes, electric eels, freshwater rays; Hanika & Kramer 1999, 2000; Stoddard 1999). The two EOD phases sum at a short distance away from the fish, attenuating the low-frequency DC component of the EOD detectible by ampullary receptors of predators. Consequently, at greater distances DC-balanced EODs are theoretically not detectible by electroreceptive predators (Stoddard 1999; Stoddard & Markham 2008; Stoddard et al. 2019). *B. bennetti* s monophasic EOD represents the asymmetrical extreme, with a large DC component and large amplitude augmenting its detectability by electroreceptive predators.

Previous studies have suggested that *B. bennetti*’s EOD functions as a Batesian mimic of the Electric Eel’s monophasic EOD (Stoddard 1999). This fish is the only gymnotiform capable of producing a strong electric discharge to incapacitate predators and prey alike. Crampton & Albert (2006) argue that this mimicry is not a likely evolutionarily stable strategy due to a large discrepancy in the relative abundance of *B. bennetti* and *E. electricus*, the former being three orders of magnitude more abundant than the latter. Instead, Crampton & Albert (2006) suggest the persistence of the monophasic EOD is related to species recognition.

Sensory bias and a female preference for signals with low frequency energy could further explain the reversion to a monophasic EOD. Sexual dimorphism has been documented in many gymnotiform species. According to Crampton & Albert (2006), *B. bennetti* males produce EODs with amplitudes 2-3 times larger than females of comparable sizes, likely because they possess relatively larger sized electrocytes compared to juvenile and females (Crampton et al. 2016b). Building on the hypothesis of compatible mate (“species”) recognition, it is possible that the monophasic EOD of *B. bennetti* could be a result of sexual selection. Consistent with handicap principle, female preference for a large amplitude could select for a costly signal both in terms of energetics and susceptibility to predation (Zahavi 1975). A mate choice study on *B. gauderio* suggests that EODa is the most salient EOD feature used by females to assess potential mates (Curtis and Stoddard 2003). Maximum EODa is largely a function of fish size, with larger individuals having more electrocytes thus producing larger EODs. Larger amplitudes also imply greater metabolic costs (larger APs require more ATP to restore electrocyte membrane potential per EOD), meaning a large amplitude could serve as an honest indicator of male condition. Still, mate choice studies with *B. bennetti* are needed to test this potential preference.

### Conclusion

This and other recent studies (Ban et al. 2015) highlight that our understanding of electrocyte physiology is still very incomplete, especially with respect to the localization of different ion channel populations on the electrocyte membrane. Researchers have recognized the importance of the heterogenous distribution of ion channels in electrocytes since early studies in the Electric Eel (Ellisman & Levinson 1982; Fritz et al. 1983), yet little is known about the regulatory mechanisms contributing to their localization. In many cases, the density and distribution of specific ion channels is critical for the function of nerve cells, such as in the nodes of Ranvier (Schulz et al. 2008) or the axon initial segment (AIS) (Kole et al. 2008), and abnormalities in channel localization can lead to neural pathologies (England et al. 1996). Further study of electrocyte ion channels in weakly electric fishes could improve our understanding of the processes governing ion channel localization as well as their evolutionary underpinnings.

In addition, these findings show the need for a better understanding of the ecology of these species and the adaptive significance of EOD plasticity. In view of *B. bennetti*’s broad geographical distribution and abundance (Crampton et al. 2016b), it seems the energetic and predation handicaps considered here have not hindered its ecological success and point to other unknown aspects of its ecology, such as mate choice preferences and levels of intraspecific competition (Gavassa et al. 2012a, b). Natural history studies of these animals are sorely missing, and sadly, we could be running out of time. Due to the metabolic demands of their unique sensory systems, weakly electric fishes may be disproportionately susceptible to environmental disturbances such as those induced by climate change and the rapid changes in land use currently occurring in the Amazon (Markham et al. 2016; Montag et al. 2019).

## Acknowledgements

We would like to thank Jose Alves Gomes for lab space and assistance in the field, Tiago Pires and Jansen Zuanon for enabling fish transport, Rosemary Knapp for cryotome use and assistance, JP Masly for use of his Zeiss microscope, and Rosalie Maltby for fish care.

